# Over 2000-fold increased production of the leaderless bacteriocin garvicin KS by genetic engineering and optimization of culture conditions

**DOI:** 10.1101/298489

**Authors:** Amar A. Telke, Kirill V. Ovchinnikov, Kiira S. Vuoristo, Geir Mathiesen, Tage Thorstensen, Dzung B. Diep

**Affiliations:** Faculty of Chemistry, Biotechnology and Food Science, Norwegian University of Life Sciences, Ås, Norway

**Author notes:** Address correspondence to Dzung B. Diep. Present address: Faculty of Chemistry, Biotechnology, and Food Science. Norwegian University of Life Sciences PB 5003 Chr. M. Falsensvei 1. N-1432 Aas Norway.

## Abstract

The leaderless bacteriocin Garvicin KS (GarKS) is a potent antimicrobial, being active against a wide range of important pathogens. GarKS production by the native producer *Lactococcus garvieae* KS1546 was however relatively low (80 BU/ml) under standard laboratory growth conditions (batch culture in GM17 at 30°C). To improve the production of GarKS, we systematically evaluated the impact of different media and media components on bacteriocin production. Based on the outcomes a new medium formulation was made to greatly improve bacteriocin production. The new medium composed of pasteurized milk and tryptone (PM-T), increased GarKS production about 60-fold compared to that achieved in GM17. GarKS production was increased further 4-fold (i.e., to 20,000 BU/ml) by increasing gene dose of the bacteriocin gene cluster (*gak*) in the native producer. Finally, a combination of the newly composed medium (PM-T), an increased gene dose and a cultivation at a constant pH 6 and a 50-60% dissolved oxygen level in growth medium, gave rise to a GarKS production of 164,000 BU/ml. This high production, which is about 2000-fold higher compared to that initially achieved in GM17, corresponds to a GarKS production of 1.2 g/L. To our knowledge, this is one of the highest bacteriocin production reported hitherto.

**Importance:** Low bacteriocin production is a well-known bottle-neck in developing bacteriocins into large-scaled and useful applications. The present study shows different approaches that significantly improve bacteriocin production. This is an important research field to better exploit the antimicrobial potential of bacteriocins, especially with regard to the decreasing effect of antibiotics in infection treatments due to the global emergence of antibiotic resistance.

## Introduction

The decreasing effectiveness of antibiotics has become a serious worldwide problem due to the emergence of multidrug-resistant bacteria (1, 2). Despite that, the number of new commercially available antibiotics is dwindling. This is partly due to the fact that developing new antibiotics is a very costly process (3), and the biopharma companies are therefore often reluctant to invest large money in new antibiotics that soon may be useless because of resistance development. Consequently, there is an urgent need of cost-effective and efficient antimicrobial agents with different killing mechanisms to overcome multidrug-resistant bacteria.

Bacteriocins are ribosomally synthesized antibacterial peptides produced by bacteria, probably as a means to compete for nutrients and habitats (4). So far hundreds of bacteriocins have been isolated and characterized. Most of them have narrow-spectrum activity but some are active against a broad-spectrum of bacteria including food-spoiling bacteria as well as important pathogens (5, 6). Bacteriocins produced by lactic acid bacteria (LAB) are particularly interesting due to LAB’s safe status as they are commonly found in our foods (7, 8) and the gastrointestinal tract of man (9) and animals (10). Most bacteriocins are membrane-active peptides, killing sensitive bacteria by membrane disruption after selective interaction with specific membrane receptors (11-15). This mode of action is different from most antibiotics which often act as enzyme-inhibitors (16, 17). For this reason, antibiotic-resistant pathogens are often sensitive to bacteriocins, thus making the latter very attractive as alternative or complementary drugs for therapeutic use, especially to fight antibiotic resistance. Nevertheless, poor production is often a bottleneck in large-scaled production of bacteriocins. Previous studies have shown that bacteriocin production can be increased by optimization of growth conditions such as cultivation temperature, pH, aeration and growth medium (18-25). In addition, various heterologous expression systems have been reported for increased bacteriocin production (26-30).

Recently we have reported the identification and characterization of a novel three-peptide bacteriocin called garvicin KS (GarKS), produced by *L. garvieae* KS1546, a strain isolated from raw bovine milk in Kosovo (31). A gene cluster (*gak*) containing the three structural genes (*gakABC*) and genes likely involved in immunity (*gakIR*) and transport (*gakT*) has been identified in the genome (31). GarKS is active against a broad spectrum of bacteria such as *Listeria, Staphylococcus*, *Bacillus*, *Streptococcus* and *Enterococcus* (31). Despite its great potential, production of GarKS is relatively moderate in standard laboratory growth conditions. To overcome this problem, we conducted a multi-factorial optimization study that resulted in over 2000-fold increased bacteriocin production. This approach includes medium optimization, genetic engineering and cultivation optimization,

## Results

### GarKS production in complex media

*L. garvieae* KS1546 (hereafter referred to KS1546) was routinely grown in the complex medium GM17 at 30°C without agitation, and GarKS production was typically of 80 BU/mL after 7-12 h growth. The bacteriocin production by KS1546 was examined in different complex media (MRS, BHI and TH). Highest production was found between 7-12 h of growth in all tested media except for TH where bacteriocin production appeared constantly low for all time-points tested (Fig. 1A). Relative to GM17, GarKS production increased 2 to 4-fold in MRS, while it was about 2 to 4-fold less in BHI and TH (Fig. 1A). Cell growth was the best in GM17 (3×10^9^ CFU/ml) but the poorest in MRS (1×10^9^ CFU/ml) after 24 h at 30°C (Table 1).

**Figure. 1.**
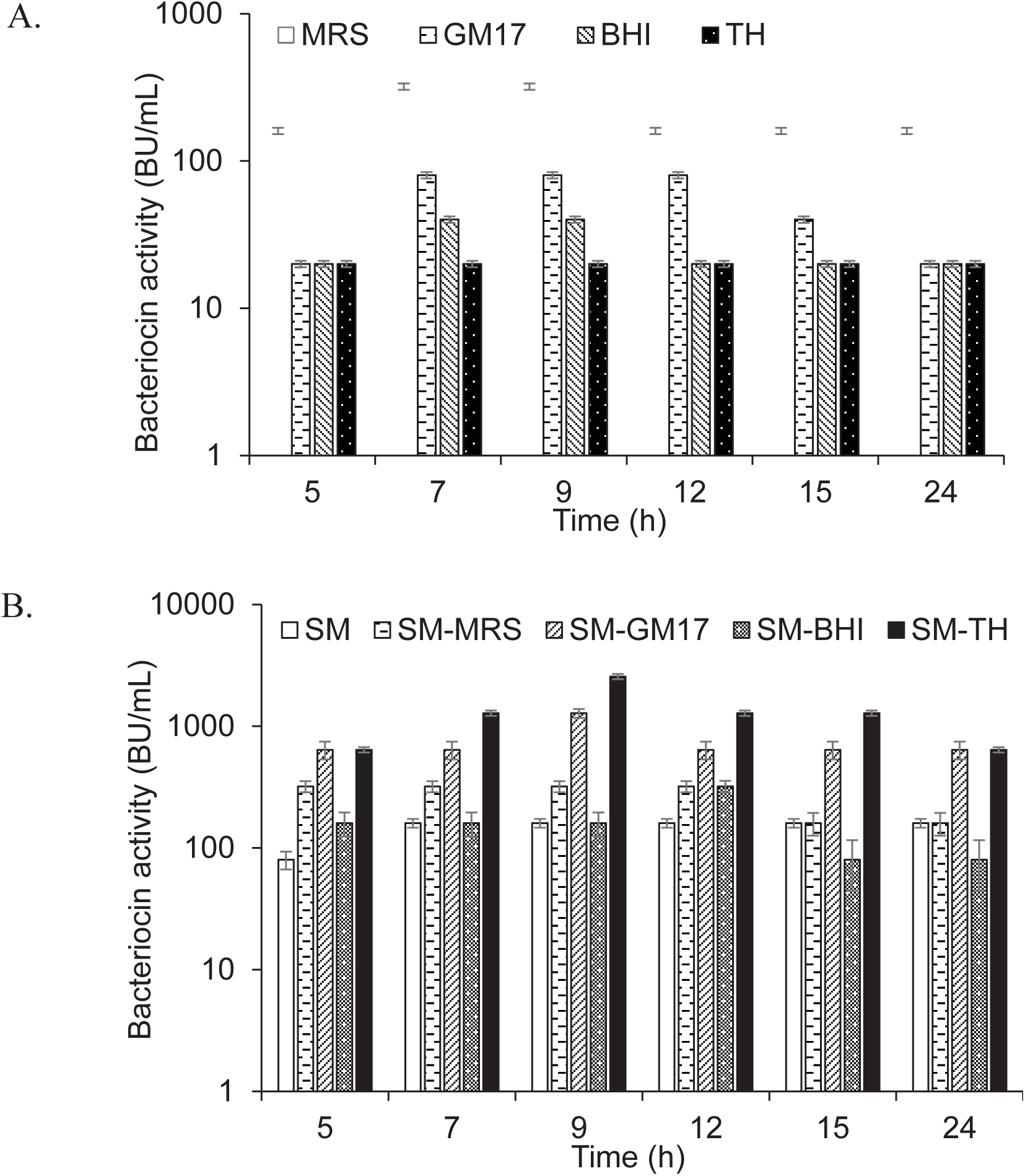
GarKS production by the native producer in different complex growth media (A), and in skim milk (SM) combined with complex growth media (B). Each culture was started by adding 1% (v/v) culture inoculum to a 5 ml growth medium and then incubated at 30°C without shaking. Bacteriocin activity was measured at different time points. Standard deviations were based on triplicate assays.

**Table 1.** Influence of growth media, increased gene dose and culture conditions on bacteriocin production.

| Strain | Growth medium | Bacteriocin activity (BU/ml) | Cell growth (CFUs/ml) |
| --- | --- | --- | --- |
| Native producer<br>*L. garvieae* KS1546 | GM17<sup>a</sup> | 80 ±20 | 3×10<sup>9</sup> |
|  | MRS<sup>a</sup> | 320 ±20 | 1×10<sup>9</sup> |
|  | BHI<sup>a</sup> | 20 | 1.5 ×10<sup>9</sup> |
|  | TH<sup>a</sup> | 20 | 2 ×10<sup>9</sup> |
|  | SM<sup>b</sup> (10%, w/v) | 160 ±20 | 2 ×10<sup>8</sup> |
|  | Tryptone<sup>a</sup> (10%, w/v) | 80 ±20 | 3 ×10<sup>8</sup> |
|  | SM-TH<sup>b</sup> | 2600 ±80 | 2.9 ×10<sup>9</sup> |
|  | SM-GM17<sup>b</sup> | 1280 | 3 ×10<sup>9</sup> |
|  | SM-MRS<sup>b</sup> | 320 | 2.8 ×10<sup>9</sup> |
|  | SM-BHI<sup>b</sup> | 160 | 2.9×10<sup>9</sup> |
|  | SM-T<sup>b</sup> | 2600 ±80 | 3 ×10<sup>9</sup> |
|  | SM-T-YE<sup>b</sup> | 1300 ±40 | 3 ×10<sup>9</sup> |
|  | PM-T<sup>b</sup> | 5100 ±80 | 3.5 ×10<sup>9</sup> |
| The recombinant<br>producer *L. garvieae*<br>KS1546-pA2T | PM-T<sup>b</sup> (uncontrolled<br>pH) | 20,000 ±400 | 3.5 ×10<sup>9</sup> |
|  | PM-T<sup>b</sup> (constant pH 5) | 2600 ±80 | 3.2 ×10<sup>9</sup> |
|  | PM-T<sup>b</sup> (constant pH 6) | 82,000 ±400 | 0.7 ×10<sup>10</sup> |
|  | PM-T<sup>b</sup> (constant pH 7) | 41,000 ±400 | 0.65 ×10<sup>10</sup> |
|  | PM-T<sup>b</sup> (constant pH 6<br>and aeration) | 164,000 ±400 | 1×10<sup>10</sup> |

The bacteriocin activity and cell growth from complex growth media (a) and milk based media (b) were measured after 7 and 9 h of incubation, respectively. SM- Skim milk, T- Tryptone, YE-Yeast extract, and PM-Pasteurized milk. Standard deviations (±) were based on triplicate assays.

### GarKS production increased in milk-based media

It is well known that bacteria are ecologically adapted to the environments where they normally thrive. Since the producer KS1546 was isolated from raw milk (31), we examine the possibility to use skim milk (SM) as growth medium. Bacteriocin production was increased 2-fold in SM (160 BU/ml) compared to GM17 (Fig. 1B). However, cell growth was remarkably poor in skim milk (2×10^8^ CFUs/ml) (Table 1), indicating that some growth factors were present in complex media but absent in SM. Therefore, we tested the mixtures (50:50; v/v) of skim milk and complex media (GM17, MRS, BHI and TH). As a result, the bacteriocin production was increased 16 times in skim milk combined with TH (SM-TH) and 8 times in SM-GM17, compared to the production in skim milk (Table 1 and Fig. 1B). The bacteriocin production in SM-TH and SM-GM17 was 2600 BU/ml and 1280 BU/ml after 9 h of incubation, respectively. On the other hand, no significant increase of GarKS in SM-MRS (320 BU/ml) and SM-BHI (160 BU/ml) was found in all time points (Fig. 1B). All medium formulations gave approximately a similar cell density, i.e., between 2.8×10^9^-3×10^9^ CFUs/ml (Table 1).

The results above indicate that bacteriocin production was significantly influenced by some specific factor(s)/nutrient(s), which are present in TH and GM17, but absent in MRS and BHI. Tryptone, a tryptic digest of milk protein casein (32), is one of the nutrients found in GM17 and TH, but not in MRS and BHI. The final concentration of tryptone in GM17 and TH broth is 0.5% and 2%, respectively. To examine whether tryptone could improve cell growth and bacteriocin production in combination with SM, we made formulations with different v/v ratios of SM and 10% tryptone (w/v). Highest bacteriocin production (about 2,600 BU/ml) was achieved when they were mixed in equal volumes (50%; v/v); this mixture had a final concentration of tryptone of 5% (w/v) (Figure 2). Under these circumstances, final cell density was comparable to that in GM17, i.e., about 3×10^9^ CFU/ml (Table 1). The formulation composed of SM (50 %; v/v) and a final 5% of tryptone (w/v) is hereafter called SM-T.

**Figure. 2.**
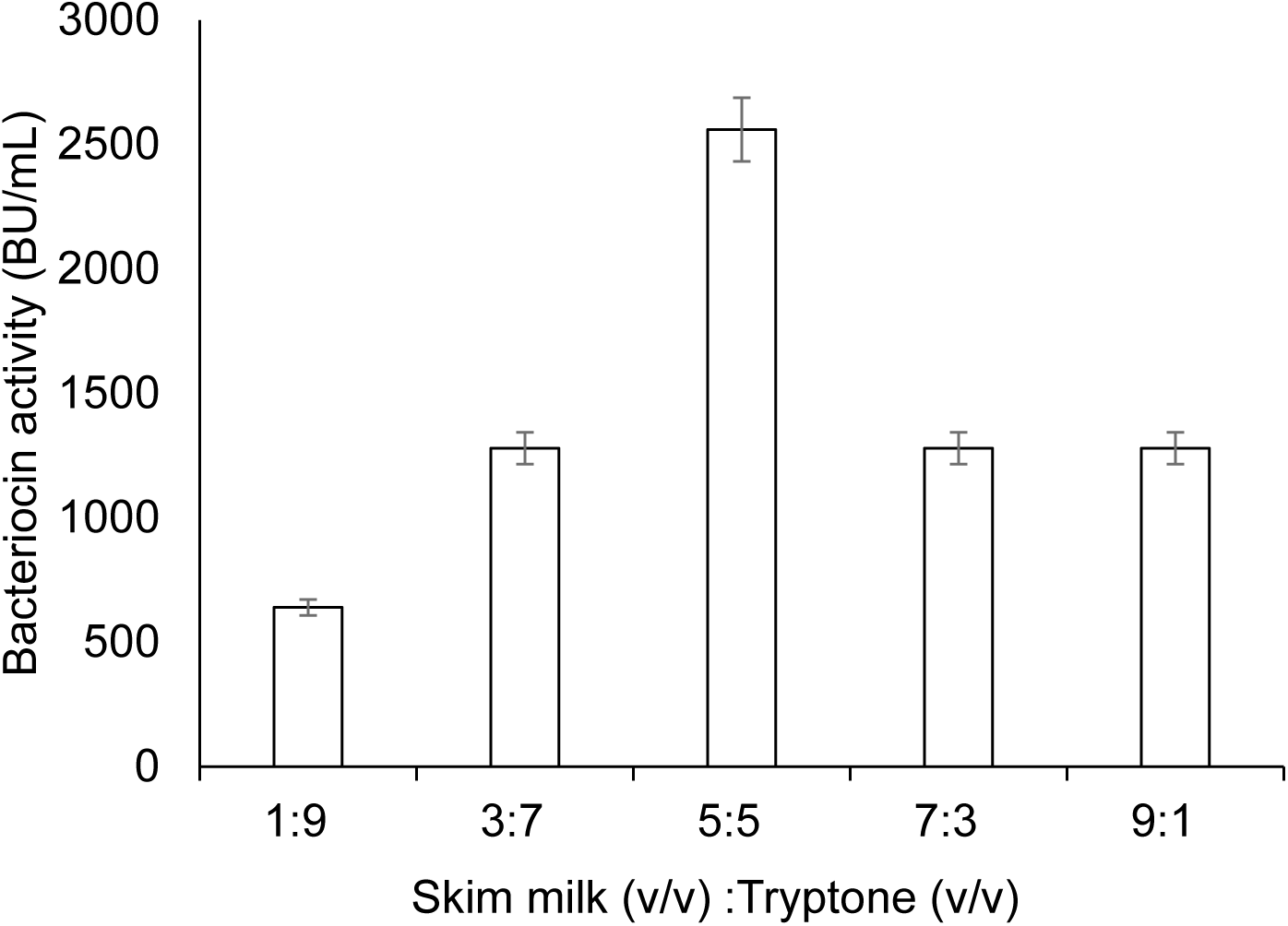
Bacteriocin production in a medium composed of skim milk and tryptone. Different ratios of skim milk and tryptone were made in the formulation by mixing an increasing portion of skim milk (10%; w/v; from 1 volume to 9 volumes) with a corresponding decreasing portion of tryptone (10%, w/v; 9 volumes to 1 volume). For growth conditions, see legend in figure 1. The bacteriocin activity was measured after 9 h of culture incubation. Standard deviations were based on triplicate assays.

Yeast extract is a rich source of vitamins, minerals, and amino acids, which often improves bacterial growth. We examined the effect of yeast extract (YE) in combination with SM-T. The resulting formulation, SM-T-YE (SM-T containing 1% (w/v) yeast extract) yielded the same cell density as in SM-T (3×10^9^ CFU/ml), but bacteriocin production was reduced by 50% (Table 1). Yeast extract was therefore excluded from the growth medium.

Although SM-T appeared as a good medium for the producer, we constantly encountered the problem associated with caramelization of milk sugars in skim milk during autoclaving, which might have detrimental effects on milk nutrition value. To avoid this problem, the autoclaved skim milk in SM-T was replaced with equal amount of pasteurized milk, resulting in a new medium termed pasteurized milk–tryptone (PM-T). The contents in milk (Q-milk) according to the manufacturer (Q-Meieriene AS, Bergen, Norway) are, g/l: fat, 5; carbohydrate, 45; protein, 35; salt, 1; calcium, 1.3; vitamin B_2_, 0.001; and vitamin B_12_, 0.7×10^-5^. Indeed, cell growth in PM-T was increased to 3.5×10^9^ CFU/ml and GarKS production increased two-fold in comparison to that in SM-T (Table 2).

**Table 2.**
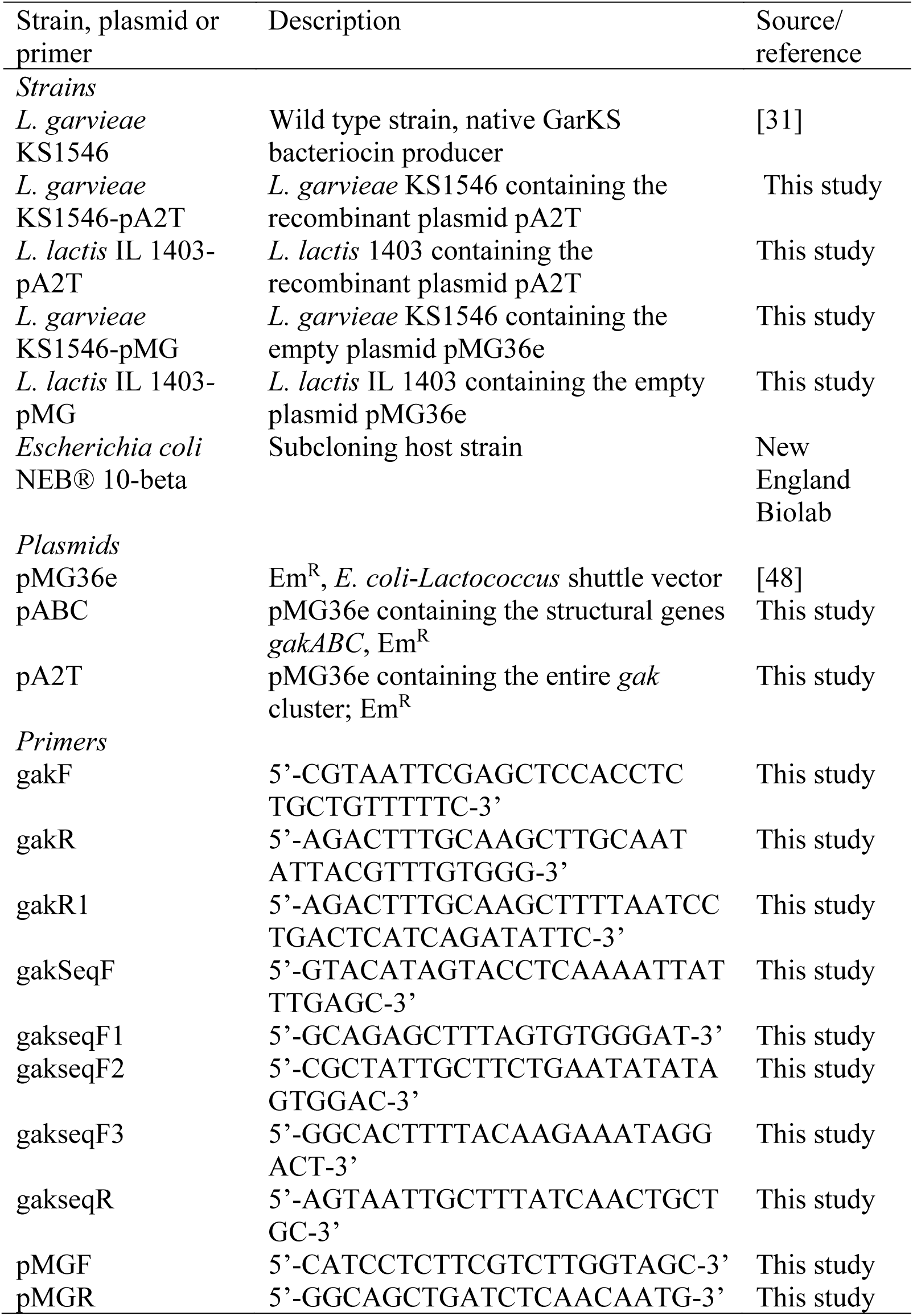
Bacterial strains, plasmids and primers used in this study

### GarKS production increased by genetic engineering

The three structural genes (*gakABC*) encoding the three peptides that constitute GarKS are clustered with genes probably involved in immunity (*gakIR*) and transport (*gakT*). First we explored the possibility to increase bacteriocin production by increasing only the gene dose of structural genes *gakABC* in the native producer. The recombinant plasmid pABC carrying structural genes *gakABC* was constructed to deliver high gene dose in the native producer (Fig. 3A). However, we failed to get any transformants even after several attempts. Similar negative result (i.e., no transformants) was obtained when we attempted to transfer pABC into the heterologous host *L. lactis* IL1403 (data not shown). Probably, increased gene dose of the structural genes might override the immunity or/and the transporter in the native producer, leading to toxicity to cells. Consequently, the plasmid pA2T carrying the entire *gak* locus including the genes involved in immunity and transport was constructed pA2T (Fig. 3B). The plasmid was first transferred into *L. lactis* IL1403. As expected, transformation was successful and bacteriocin production was detected in transformed cells (data not shown), confirming the functionality of the *gak* locus. Next the plasmid was transferred into the native KS1546 and the clone (KS1546-PA2T) was assessed for bacteriocin production. Using PM-T as growth medium, GarKS production by the recombinant producer KS1546-pA2T was found to increase to 20,000 BU/mL, which is about 4 times more than the production without increased gene dose (native KS1546 in PM-T), and about 250-fold more than that initially obtained in GM17 (native KS1546 in GM17) (Table 1). To compare the growth patterns, the native and recombinant producers were grown under similar growth conditions. The recombinant producer KS1546-pA2T showed a prolonged lag growth phase compared to the native GarKS producer (with or without empty plasmid). Nevertheless, KS1546-pA2T reached eventually about the same high cell density as the wild type control cells when it entered stationary growth phase (see Fig.4).

**Figure. 3.**
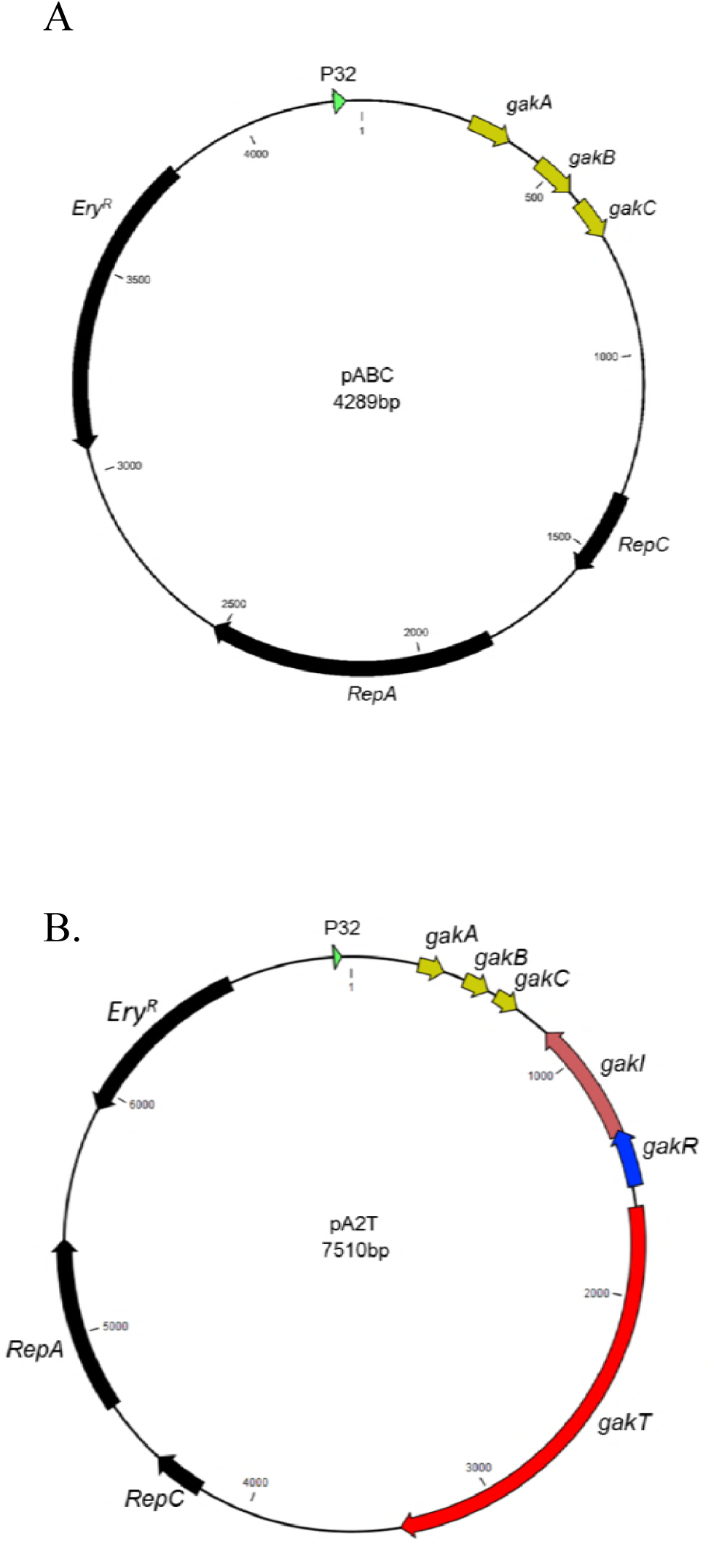
The plasmid map of pABC (A) and pA2T (B), which were used to increase the gene dose of the structural genes (*gakABC*) and the *gak* cluster in the native producer, respectively.

**Figure. 4.**
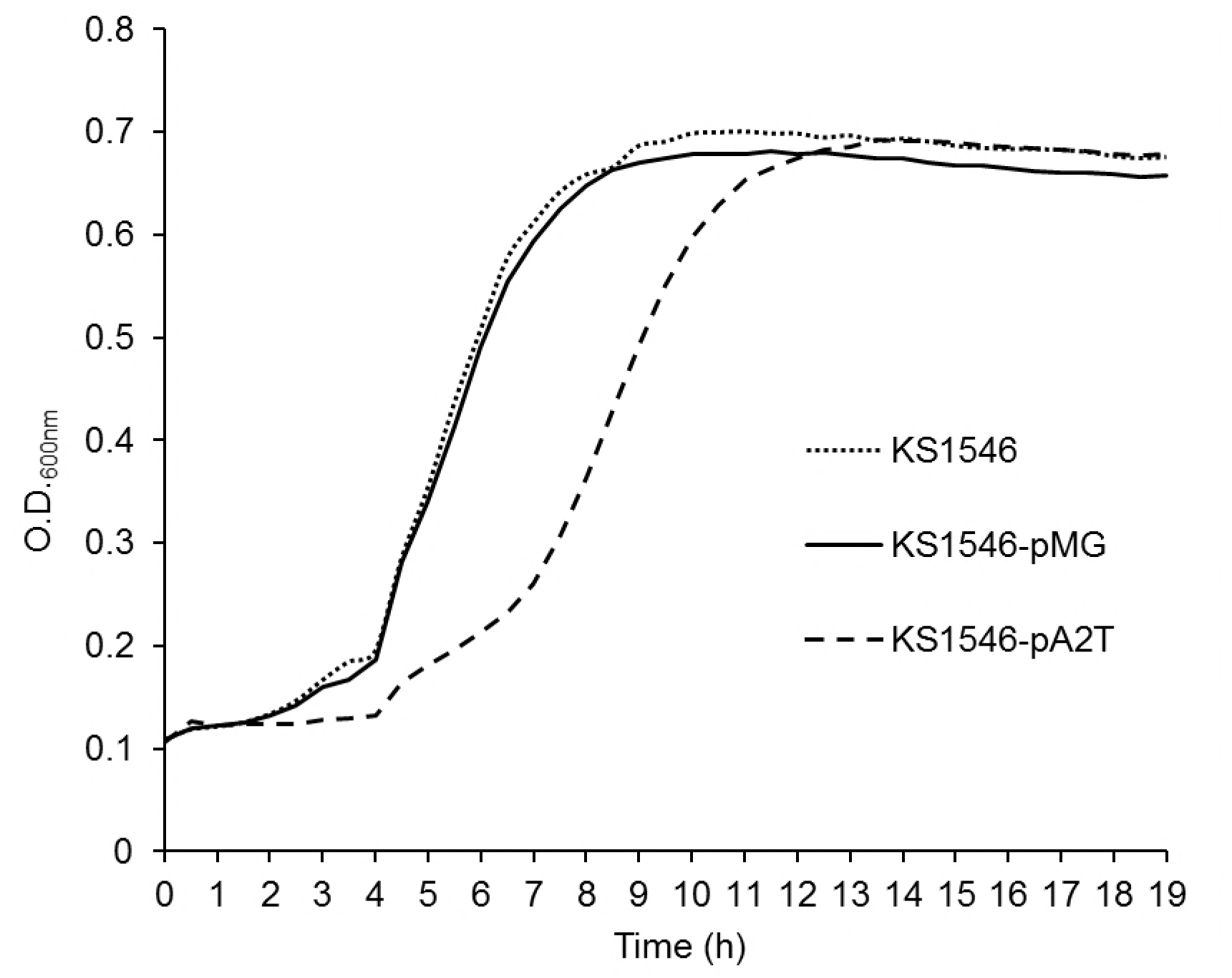
Temporal growth profile of the recombinant producer (KS1546-pA2T), and the native producer with empty plasmid (KS1546-pMG) or without plasmid (*L. garvieae* KS1546). Data were acquired from triplicate assays. Standard deviations are within a range ±0.01 to ±0.05.

### Optimization of culture conditions in a bioreactor increased GarKS production

The initial pH at 7 was declined to 4.8 when the recombinant producer KS1546-pA2T was grown in PM-T for 6-7 h at 30°C (data not shown). To examine whether pH reduction could have a negative impact on bacteriocin production, we grew the recombinant producer (KS1546-pA2T) in PM-T in a bioreactor with constant pH at 5, 6 or 7. Indeed, pH had a great impact on cell growth and bacteriocin production. Highest cell growth (0.7 ×10^10^ CFU/ml) and bacteriocin production (82,000 BU/ml) were found at constant pH 6 (Table 1). Bacteriocin production measured at all time-points was also highest at constant pH 6 (Fig. 5). Cell growth and bacteriocin production were lowest at constant pH 5.

**Figure. 5.**
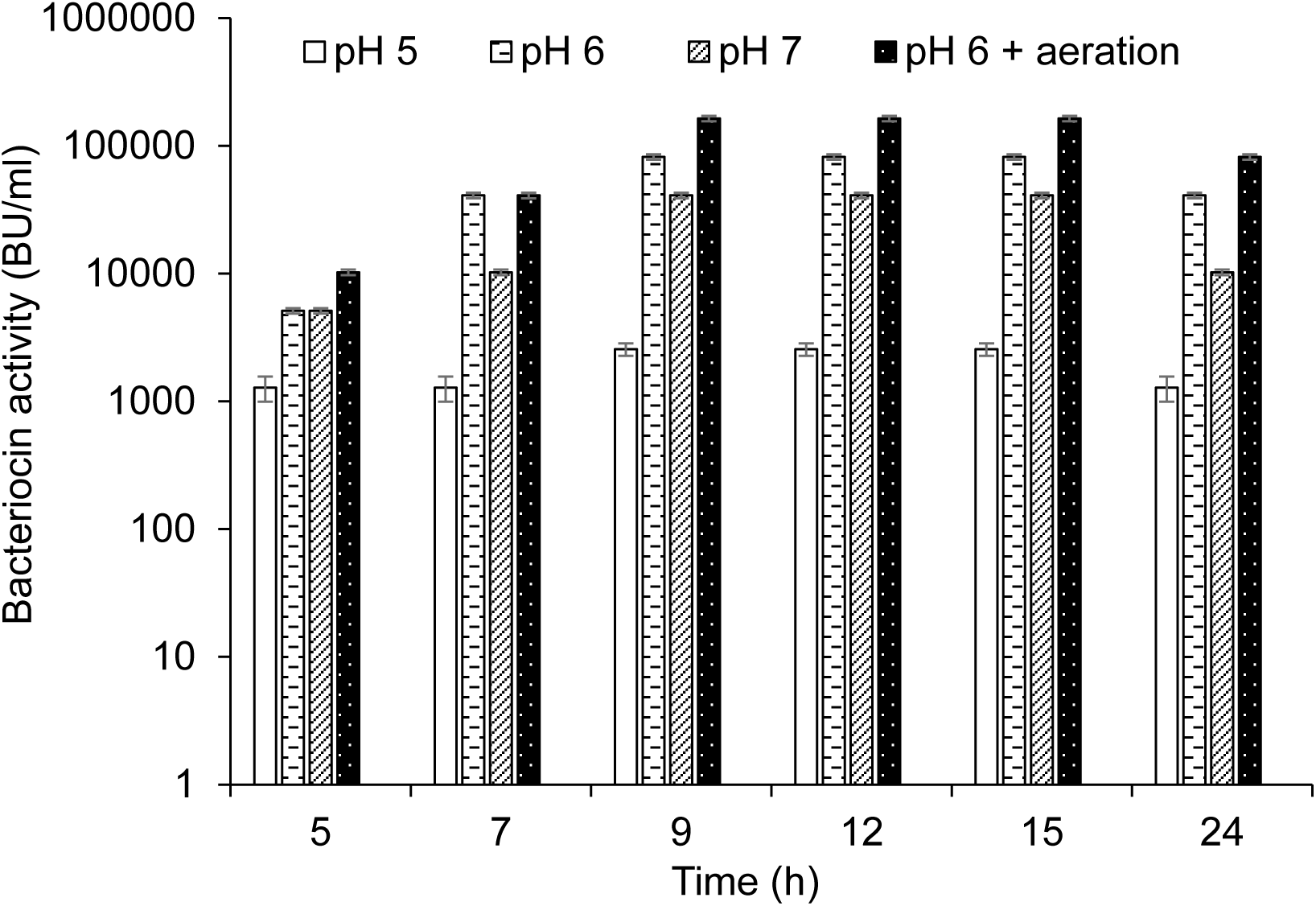
GarKS production of the recombinant producer (KS1546-pA2T) in cultivation at constant pH (pH at 5, 6, or 7) or at constant pH 6 and aeration (50-60 % dissolved oxygen). Each culture was started by adding 2% (v/v) culture inoculum in 1.5 l of PM-T medium containing erythromycin at final concentration of 5 µg/ml. Standard deviations were based on triplicate assays.

Aeration is defined as dissolved oxygen (DO) percentage in a culture medium. We observed that the initial DO level at 50-60% was declined to 10% after 2 hours of cell growth in PM-T medium and at constant pH 6. The effect of aeration on GarKS production was therefore examined by purging the atmospheric sterile air into the growth medium (constant pH at 6). With aeration kept at 50-60%, highest cell growth (1×10^10^ CFU/ml) and bacteriocin production (164,000 BU/ml) were obtained (Table 1 and Fig. 5). This level of bacteriocin production (164,000 BU/ml) was about 2000-fold more than the initial production in GM17 which was 80 BU/ml.

We have previously shown that synthetic GarKS is functionally comparable to the biologically produced counterpart (31). Synthetic GarKS (with > 90% purity) has a specific activity of 130-140 BU/µg. Hence, the production of 164,000 BU/ml is equivalent to 1.2 g GarKS per liter which is a level of commercial importance.

## Discussion

GarKS is potent against a set of important pathogens including *Staphylococcus*, *Bacillus*, *Listeria, Streptococcus* and *Enterococcus*, making it very attractive in diverse antimicrobial applications from food to medicine. Unfortunately, as also for many other bacteriocins, GarKS is produced at relatively low levels during normal laboratory growth conditions (31). The low production by the native producer can dramatically hamper potential applications of GarKS as industrial use of bacteriocins requires high and cost-effective production. We have shown that optimization of bacteriocin production by a bacterial strain is multi-factorial process, which involves the systematic evaluation of nutritional ingredients and growth conditions e.g., temperature, pH and aeration. The type of growth medium is probably one of the key factors in bacteriocins production (33). The complex media e.g., GM17, MRS, BHI, and TH have been used in cultivation of LAB because they give relatively good cell growth in laboratory conditions but not necessary for bacteriocin production (34). This is illustrated in our study, GarKS production was the best in MRS (320 BU/ml) but the lowest in BHI and TH (both 20 BU/ml) while the cell growth appeared about in the same range in these media (1-2×10^9^ CFU/ml).

To choose the optimal medium for bacteriocin production is often an empirical matter. The components from complex media influencing bacteriocin production are often elusive and the outcomes might vary significantly dependent on the type of producers. Nevertheless, some media components have been shown to enhance bacteriocin production by inducing stress conditions due to nutrient limitation (35) or stabilizing the bacteriocin molecules (36). The use of commercial complex media (e.g., MRS) is not a cost-effective approach for large-scale bacteriocin production. For instance, culture medium could account for up to 30% of the total production cost in commercial biomolecule production (37). Accordingly, high costs of complex media will reduce attractiveness of bacteriocins for commercial application. Our bacteriocin producer is a strain of *L. garvieae* isolated from raw milk and it has the capacity to ferment milk-associated sugars such as lactose and galactose while another strain of *L. garvieae* isolated from intestine of Mallard duck can not (31). Milk is a low-cost product relative to complex media and could be an ideal medium for GarKS producer. However, the native producer appeared to grow poorly in sole skim milk that might be the reason for reduced GarKS production. Skim milk is enriched in lactose and galactose as carbon source but does not contain easily accessed nitrogen-containing components for bacteria. Thus, the combination of tryptone and pasteurized skim milk which was found best for cell growth was in line with the notion that tryptone serves as an enriched source of nitrogen. Further, this formula also increased bacteriocin production over 30 fold compared to the growth in GM17. Tryptone is composed of short peptides that are derived from enzymatic digest of milk protein casein and serves as an enriched source of nitrogen in bacterial growth media.

Increase of gene dose is another means to enhance the production of biomolecules (38). In the present study, we observed a 4-fold increase in bacteriocin production when a plasmid carrying the entire *gak* locus was introduced into the native producer. Interestingly, when we attempted to increase gene dose by introducing the structural genes only (using the plasmid pABC), no transformed cells were obtained. One possible explanation for this negative outcome is that expression of genetic determinants involved in bacteriocin iosynthesis is often highly fine-tuned to secure immunity and efficient export. The extra gene dose of the structural genes alone might override either immunity and/or transporter proteins, leading to toxicity in cell and cell death. It is worth mentioning that most bacteriocins are expressed with a leader sequence which is necessary not only for export but also to keep the bacteriocins in inactive form before export. For leaderless bacteriocins like in the case of GarKS, they are produced in mature active forms before export, therefore an intracellular dedicated protection mechanism (immunity) available is crucial for cell survival.

We and others have observed that bacteriocin production by a certain strain is unstable, and dependent on the culture conditions applied (39, 40). Consequently, different growth parameters were examined to optimize the production of GarKS. LABs are well known for reducing culture pH due to lactic acid production (41) and this is also true for the GarKS producer. We found that culture conditions with constant pH 6 favors the cell growth and a high level of GarKS production. Similarly, optimal nisin production has been reported at constant pH 6.5 (42). The availability of oxygen also has a great influence on microbial cell growth and metabolic activities (43). Microorganisms vary with respect to their requirements and tolerance toward molecular oxygen. *L. garvieae* is a facultative anaerobic microorganism and its metabolic activities have been reported to differ between aerobic and anaerobic conditions (44). We observed that the controlled aeration had a positive effect on the cell growth and bacteriocin production. Similar results have also been observed for other bacteriocins. For example, nisin A production by *L. lactis* UL719 was enhanced with aeration (45). On the other hand, aeration has also been reported to be antagonistic to the production of lactosin S (46) and LIQ-4 bacteriocin (47), suggesting that the effect of aeration on bacteriocin production is strain-dependent.

In terms of cost-effectiveness, the medium PM-T contained tryptone which is a relatively costly component; therefore we are searching for alternatives to replace tryptone. In preliminary studies, we tested the chicken hydrolysate (processed from a waste product from meat industry) as an alternative low-cost protein source to produce GarKS. We found that the recombinant producer grew well in a medium based on Pasteurized milk and chicken hydrolysate (PM-CH), yielding a cell density of 3×10^9^ CFU/ml. However, although GarKS production in PM-CH was 8 times better than in the complex media GM17, the production was 8 times less than in PM-T. Thus, further studies are necessary to optimize a PM-CH-based medium in order to achieve high level and cost-effective bacteriocin production.

Low bacteriocin production is often a bottle-neck in large-scaled production of bacteriocins for commercial use. Optimization of bacteriocin production is therefore an important research field to better exploit the antimicrobial potential of bacteriocins, especially with regard to the decreasing effects of antibiotics in infection treatments due to the global emergence of antibiotic resistance. In the present study we have shown that we managed to achieve a very high level of GarKS production, amounting to 164,000 BU/ml, by combining medium optimization, genetic engineering and culture condition optimization. This amount is about 2,000 times higher compared to the initial production in GM17 (80 BU/ml). A production of 164,000 BU/ml is equivalent to 1.2 g GarKS per liter. To our knowledge, this is one of the highest bacteriocin production achieved so far. In comparison, nisin production has been reported to 0.40-0.80 g/L by *L. lactis* grown in a medium composed of equal volume of skim milk and complex media GM17 (5). Finally, our study and others’ have shown that optimization of bacteriocin production is an empirical and multi-factorial process and that it is highly strain-dependent. Only by systematic evaluation of different aspects influencing growth and gene regulation one can find conditions suitable for high levels of production.

## Materials and Methods

### Bacterial strains and growth conditions

All bacterial strains and plasmids used in this study are listed in Table 2. Unless otherwise stated, the native bacteriocin producer *L. garvieae* KS1546 was grown in M17 broth supplemented with 0.5% glucose (GM17) under static condition at 30°C. NEB® 10-beta *E. coli* (New England Biolabs, Beverly, MA, USA) was grown in Luria-Bertani (LB) broth with shaking (200 rpm) at 37°C. Bacterial culture media and supplements were obtained from Oxoid Ltd (Hampshire, UK). When necessary, erythromycin (Sigma-Aldrich Inc., St. Louis, MO, USA) was added at 200 µg/ml for *E. coli* and at 5 µg/ml for lactococcal species.

### Growth media for GarKS production

The influence of different growth media on GarKS production was assessed in batch cultures under static condition at 30°C. Following commercial complex media were used: GM17, deMan, Rogosa and Sharpe (MRS), Todd-Hewitt (TH) and Brain Heart Infusion (BHI). To make new milk-based medium formulations, skim milk (5%, w/v) or pasteurized skim milk was combined with an equal volume of GM17, MRS, TH, and BHI, or with tryptone (10% w/v). Skim milk (SM) was prepared by using milk powder (Oxoid, UK) while pasteurized milk (PM) was obtained from a dairy company in Norway, Q-milk.

### DNA manipulation

The *gak* cluster responsible for production of GarKS was amplified from genomic DNA of *L. garvieae* KS1546 using Phusion High-fidelity DNA polymerase (New England Biolabs, UK) and the primers gakF and gakR1 (Table 2). The genes *gakABC* encoding the three peptides constituting GarKS were amplified using the primers gakF and gakR (Table 2). Restriction sites SacI and HindIII were introduced at the 5’end of forward and reverse primers. NEBuilder HiFi DNA assembly cloning kit (New England Biolabs) was used to assemble the PCR fragments into the plasmid pMG36e (48). Plasmid DNA was amplified in *E. coli* NEB® 10-beta before being transferred into *L. garvieae* KS1546 or *L. lactis* IL1403 cells using a Gene Pulser^TM^ (Bio-Rad Laboratories, Hercules, CA, USA). Primers used in this study were obtained from Life Technologies AS (Thermofisher Scientific, Oslo, Norway). The integrity of all recombinant plasmids was confirmed by Sanger DNA sequencing (GATC Biotech AG; Constance, Germany), which were sequenced using primers gakseqF, gakseqF1, gakseqF2, gakseqF3, gakseqR, pMGF and pMGR (Table 2).

### Optimization of bacteriocin production in bioreactor conditions

The effects of pH and aeration on GarKS production were tested at various constant pH (5, 6 and 7), and at controlled aeration in a fully automated 2.5 L miniforce bioreactor (Infors AG, Switzerland). The pH was controlled by automatic addition of 5 M HCl or 5 M NaOH. The aeration was maintained by purging sterile air into culture medium. Temperature (30°C) and agitation speed of 150 rpm were maintained constant for all experiments. Samples of 2 ml were withdrawn aseptically every 2 h for determination of bacteriocin production and cell growth (see below).

### Determination of bacteriocin production and cell growth

Bacteriocin activity was measured from heat-inactivated (100°C for 10 min) cell-free culture supernatants. Bacteriocin activity was quantified using a microtiter plate assay as previously described (27, 31). One bacteriocin unit (BU) was defined as the minimum amount of the bacteriocin that inhibited at least 50% of growth of the indicator (*L. lactis* IL103) in a 200 µl culture volume. Growth curve was determined by measuring turbidity of culture at OD_600_ every 30 min for 24 h or by counting colony forming units (CFU) from serially diluted bacterial cultures on agar plates. A standard curve based on the activity of 98% pure synthetic GarKS peptides (Pepmic Co., LTD, China) was used to define the specific bacteriocin activity (BU/mg) from cell free supernatant.

## ACKNOWLEDGEMENTS

We thank Linda Godager and Line Degn Hansen for technical assistance. This study was financed by the Research Council of Norway, Project nr 254784.

## REFERENCES

1. Bush K, Courvalin P, Dantas G, Davies J, Eisenstein B, Huovinen P, Jacoby GA, Kishony R, Kreiswirth BN, Kutter E, Lerner SA, Levy S, Lewis K, Lomovskaya O, Miller JH, Mobashery S, Piddock LJ, Projan S, Thomas CM, Tomasz A, Tulkens PM, Walsh TR, Watson JD, Witkowski J, Witte W, Wright G, Yeh P, Zgurskaya HI. 2011. Tackling antibiotic resistance. Nat Rev Microbiol 9:894–6.

2. Laxminarayan R, Matsoso P, Pant S, Brower C, Rottingen JA, Klugman K, Davies S. 2016. Access to effective antimicrobials: a worldwide challenge. Lancet 387:168–75.

3. Holmes P, Mauer J. 2016. Antimicrobial Resistance And New Antibiotics. Health Aff (Millwood) 35:1935.

4. Cotter PD, Hill C, Ross RP. 2005. Bacteriocins: developing innate immunity for food. Nat Rev Microbiol 3:777–88.

5. de Arauz LJ, Jozala AF, Mazzola PG, Penna TCV. 2009. Nisin biotechnological production and application: a review. Trends in Food Science & Technology 20:146–154.

6. Chikindas ML, Garcia-Garcera MJ, Driessen AJ, Ledeboer AM, Nissen-Meyer J, Nes IF, Abee T, Konings WN, Venema G. 1993. Pediocin PA-1, a bacteriocin from Pediococcus acidilactici PAC1.0, forms hydrophilic pores in the cytoplasmic membrane of target cells. Appl Environ Microbiol 59:3577–84.

7. Henning C, Vijayakumar P, Adhikari R, Jagannathan B, Gautam D, Muriana PM. 2015. Isolation and Taxonomic Identity of Bacteriocin-Producing Lactic Acid Bacteria from Retail Foods and Animal Sources. Microorganisms 3:80–93.

8. Grosu-Tudor SS, Stancu MM, Pelinescu D, Zamfir M. 2014. Characterization of some bacteriocins produced by lactic acid bacteria isolated from fermented foods. World Journal of Microbiology & Biotechnology 30:2459–2469.

9. Millette M, Dupont C, Shareck F, Ruiz MT, Archambault D, Lacroix M. 2008. Purification and identification of the pediocin produced by Pediococcus acidilactici MM33, a new human intestinal strain. J Appl Microbiol 104:269–75.

10. O’Shea EF, Gardiner GE, O’Connor PM, Mills S, Ross RP, Hill C. 2009. Characterization of enterocin-and salivaricin-producing lactic acid bacteria from the mammalian gastrointestinal tract. FEMS Microbiol Lett 291:24–34.

11. Hasper HE, Kramer NE, Smith JL, Hillman JD, Zachariah C, Kuipers OP, de Kruijff B, Breukink E. 2006. An alternative bactericidal mechanism of action for lantibiotic peptides that target lipid II. Science 313:1636–1637.

12. Diep DB, Skaugen M, Salehian Z, Holo H, Nes IF. 2007. Common mechanisms of target cell recognition and immunity for class II bacteriocins. Proc Natl Acad Sci U S A 104:2384–9.

13. Nissen-Meyer J, Rogne P, Oppegard C, Haugen HS, Kristiansen PE. 2009. Structure-function relationships of the non-lanthionine-containing peptide (class II) bacteriocins produced by gram-positive bacteria. Curr Pharm Biotechnol 10:19–37.

14. Oscariz JC, Pisabarro AG. 2001. Classification and mode of action of membrane-active bacteriocins produced by gram-positive bacteria. Int Microbiol 4:13–9.

15. Tymoszewska A, Diep DB, Wirtek P, Aleksandrzak-Piekarczyk T. 2017. The Non-Lantibiotic Bacteriocin Garvicin Q Targets Man-PTS in a Broad Spectrum of Sensitive Bacterial Genera. Sci Rep 7:8359.

16. Davis BD. 1987. Mechanism of bactericidal action of aminoglycosides. Microbiol Rev 51:341–50.

17. Bush K, Jacoby GA. 2010. Updated functional classification of beta-lactamases. Antimicrob Agents Chemother 54:969–76.

18. Penna TCV, Moraes DA. 2002. Optimization of nisin production by Lactococcus lactis. Applied Biochemistry and Biotechnology 98:775–789.

19. Guerra NP, Pastrana L. 2002. Nisin and pediocin production on mussel-processing waste supplemented with glucose and five nitrogen sources. Letters in Applied Microbiology 34:114–118.

20. Cabo ML, Murado MA, Gonzalez P, Pastoriza L. 2001. Effects of aeration and pH gradient on nisin production. A mathematical model. Enzyme and Microbial Technology 29:264–273.

21. Tafreshi SYH, Mirdamadi S, Norouzian D, Khatami S, Sardari S. 2010. Optimization of Non-Nutritional Factors for a Cost-Effective Enhancement of Nisin Production Using Orthogonal Array Method. Probiotics and Antimicrobial Proteins 2:267–273.

22. Biswas SR, Ray P, Johnson MC, Ray B. 1991. Influence of Growth-Conditions on the Production of a Bacteriocin, Pediocin Ach, by Pediococcus-Acidilactici H. Applied and Environmental Microbiology 57:1265–1267.

23. Nel HA, Bauer R, Vandamme EJ, Dicks LMT. 2001. Growth optimization of Pediococcus damnosus NCFB 1832 and the influence of pH and nutrients on the production of pediocin PD-1. Journal of Applied Microbiology 91:1131–1138.

24. Parente E, Ricciardi A. 1994. Influence of Ph on the Production of Enterocin-1146 during Batch Fermentation. Letters in Applied Microbiology 19:12–15.

25. Aasen IM, Moretro T, Katla T, Axelsson L, Storro I. 2000. Influence of complex nutrients, temperature and pH on bacteriocin production by Lactobacillus sakei CCUG 42687. Applied Microbiology and Biotechnology 53:159–166.

26. Mesa-Pereira B, O’Connor PM, Rea MC, Cotter PD, Hill C, Ross RP. 2017. Controlled functional expression of the bacteriocins pediocin PA-1 and bactofencin A in Escherichia coli. Sci Rep 7:3069.

27. Jimenez JJ, Diep DB, Borrero J, Gutiez L, Arbulu S, Nes IF, Herranz C, Cintas LM, Hernandez PE. 2015. Cloning strategies for heterologous expression of the bacteriocin enterocin A by Lactobacillus sakei Lb790, Lb. plantarum NC8 and Lb. casei CECT475. Microbial Cell Factories 14.

28. Horn N, Fernandez A, Dodd HM, Gasson MJ, Rodriguez JM. 2004. Nisin-controlled production of pediocin PA-1 and colicin V in nisin- and non-nisin-producing Lactococcus lactis strains. Appl Environ Microbiol 70:5030–2.

29. Jiang H, Li P, Gu Q. 2016. Heterologous expression and purification of plantaricin NC8, a two-peptide bacteriocin against Salmonella spp. from Lactobacillus plantarum ZJ316. Protein Expr Purif 127:28–34.

30. Kong W, Lu T. 2014. Cloning and optimization of a nisin biosynthesis pathway for bacteriocin harvest. ACS Synth Biol 3:439–45.

31. Ovchinnikov KV, Chi H, Mehmeti I, Holo H, Nes IF, Diep DB. 2016. Novel Group of Leaderless Multipeptide Bacteriocins from Gram-Positive Bacteria. Applied and Environmental Microbiology 82:5216–5224.

32. Oh S, Rheem S, Sim J, Kim S, Baek Y. 1995. Optimizing Conditions for the Growth of Lactobacillus-Casei Yit-9018 in Tryptone-Yeast Extract-Glucose Medium by Using Response-Surface Methodology. Applied and Environmental Microbiology 61:3809–3814.

33. Guerra NP, Agrasar AT, Macias CL, Pastrana L. 2005. Modelling the fed-batch production of pediocin using mussel processing wastes. Process Biochemistry 40:1071–1083.

34. Mataragas M, Drosinos EH, Tsakalidou E, Metaxopoulos J. 2004. Influence of nutrients on growth and bacteriocin production by Leuconostoc mesenteroides L124 and Lactobacillus curvatus L442. Antonie Van Leeuwenhoek International Journal of General and Molecular Microbiology 85:191–198.

35. Verluyten J, Leroy F, de Vuyst L. 2004. Influence of complex nutrient source on growth of and curvacin A production by sausage isolate Lactobacillus curvatus LTH 1174. Applied and Environmental Microbiology 70:5081–5088.

36. Herranz C, Martinez JM, Rodriguez JM, Hernandez PE, Cintas LM. 2001. Optimization of enterocin P production by batch fermentation of Enterococcus faecium P13 at constant pH. Applied Microbiology and Biotechnology 56:378–383.

37. Rivas B, Moldes AB, Dominguez JM, Parajo JC. 2004. Development of culture media containing spent yeast cells of Debaryomyces hansenii and corn steep liquor for lactic acid production with Lactobacillus rhamnosus. International Journal of Food Microbiology 97:93–98.

38. Nijland JG, Ebbendorf B, Woszczynska M, Boer R, Bovenberg RAL, Driessen AJM. 2010. Nonlinear Biosynthetic Gene Cluster Dose Effect on Penicillin Production by Penicillium chrysogenum. Applied and Environmental Microbiology 76:7109–7115.

39. Criado R, Gutierrez J, Martin M, Herranz C, Hernandez PE, Cintas LM. 2006. Immunochemical characterization of temperature-regulated production of enterocin L50 (EntL50A and EntL50B), enterocin P, and enterocin Q by Enterococcus faecium L50. Appl Environ Microbiol 72:7634–43.

40. Diep DB, Axelsson L, Grefsli C, Nes IF. 2000. The synthesis of the bacteriocin sakacin A is a temperature-sensitive process regulated by a pheromone peptide through a three-component regulatory system. Microbiology-Uk 146:2155–2160.

41. Bartkiene E, Krungleviciute V, Juodeikiene G, Vidmantiene D, Maknickiene Z. 2015. Solid state fermentation with lactic acid bacteria to improve the nutritional quality of lupin and soya bean. Journal of the Science of Food and Agriculture 95:1336–1342.

42. Gonzalez-Toledo SY, Dominguez-Dominguez J, Garcia-Almendarez BE, Prado-Barragan LA, Regalado-Gonzalez C. 2010. Optimization of Nisin Production by Lactococcus lactis UQ2 Using Supplemented Whey as Alternative Culture Medium. Journal of Food Science 75:M347–M353.

43. Garcia-Ochoa F, Gomez E. 2009. Bioreactor scale-up and oxygen transfer rate in microbial processes: an overview. Biotechnol Adv 27:153–76.

44. Delpech P, Rifa E, Ball G, Nidelet S, Dubois E, Gagne G, Montel MC, Delbes C, Bornes S. 2017. New Insights into the Anti-pathogenic Potential of Lactococcus garvieae against Staphylococcus aureus Based on RNA Sequencing Profiling. Frontiers in Microbiology 8.

45. Amiali MN, Lacroix C, Simard RE. 1998. High nisin Z production by Lactococcus lactis UL719 in whey permeate with aeration. World Journal of Microbiology & Biotechnology 14:887–894.

46. Mortvedtabildgaard CI, Nissenmeyer J, Jelle B, Grenov B, Skaugen M, Nes IF. 1995. Production and Ph-Dependent Bactericidal Activity of Lactocin-S, a Lantibiotic from Lactobacillus-Sake-L45. Applied and Environmental Microbiology 61:175–179.

47. Kuhnen E, Sahl HG, Brandis H. 1985. Purification and Properties of Liq 4, an Antibacterial Substance Produced by Streptococcus-Faecalis Var Liquefaciens-K4. Journal of General Microbiology 131:1925–1932.

48. van de Guchte M, van der Vossen JM, Kok J, Venema G. 1989. Construction of a lactococcal expression vector: expression of hen egg white lysozyme in Lactococcus lactis subsp. lactis. Appl Environ Microbiol 55:224–8.

